# Structural brain alterations in youth with psychosis and bipolar spectrum symptoms

**DOI:** 10.1101/427609

**Authors:** Maria Jalbrzikowski, David Freedman, Catherine E. Hegarty, Eva Mennigen, Katherine H. Karlsgodt, Loes M. Olde Loohuis, Roel A. Ophoff, Raquel E. Gur, Carrie E. Bearden

## Abstract

**Objective:** Adults with established diagnoses of serious mental illness (bipolar disorder and schizophrenia) exhibit structural brain abnormalities, yet less is known about how such abnormalities manifest earlier in development.

**Methods:** We analyzed the data publicly available from the Philadelphia Neurodevelopmental Cohort (PNC). Structural magnetic resonance neuroimaging data (sMRI) were collected on a subset of the PNC (N=989, ages 9-22 years old). We calculated measures of cortical thickness (CT) and surface area (SA), along with subcortical volumes. Study participants were assessed for psychiatric symptomatology via structured interview and the following groups were created: typically developing (TD, N=376), psychosis spectrum (PS, N=113), bipolar spectrum (BP, N=117), and BP + PS (N=109). We examined group and developmental differences in sMRI measures. We also examined to what extent any structural aberration was related to neurocognition, global functioning, and clinical symptomatology.

**Results:** In comparison to all other groups, PS youth exhibited significantly reduced SA in orbitofrontal, cingulate, precentral, and postcentral regions. PS youth also exhibited reduced thalamic volume in comparison to all other groups. Strongest effects for precentral and posterior cingulate SA reductions were seen during early adolescence (ages 13-15) in PS youth. Strongest effects for reductions in thalamic volume and orbitofrontal and postcentral SA were observed in mid-adolescence (16-18 years) in PS youth. Across groups, better overall functioning was associated with increased lateral orbitofrontal SA. Increased postcentral SA was associated with better executive cognition and less severe negative symptoms in the entire sample.

**Conclusion:** In a community-based sample, we found that reduced cortical SA and thalamic volume are present early in adolescent development in youth with psychosis spectrum symptoms, but not in youth with bipolar spectrum symptoms, or with both bipolar and psychosis spectrum symptoms. These findings point to potential biological distinctions between psychosis and bipolar spectrum conditions, which may suggest additional biomarkers relevant to early identification.

## Introduction

The transition from adolescence to adulthood is a unique period of development supported by specialized brain maturation, and is also a time when the prevalence of psychiatric disorders markedly increases^1^. Severe psychiatric disorders, such as bipolar disorder and schizophrenia, likely arise via deviations from typical neurodevelopmental trajectories, although the exact nature of these changes remains unknown. While structural brain alterations in adults with bipolar disorder and psychosis are well established^2-7^, less is known about the manner in which these alterations emerge, and whether youth experiencing a broad spectrum of psychosis- and bipolar-associated symptomatology exhibit similar alterations.

Importantly, structural brain volume can be decomposed into cortical thickness (CT) and surface area (SA) which may be driven by distinct genetic and neurobiological mechanisms^8-10^. Recent large scale meta-analyses have found that, in comparison to healthy controls, there are widespread SA deficits across the cortex in adult patients with schizophrenia^4^, which were not observed in a large-scale meta-analysis of cortical structure in bipolar disorder^5^. This pattern was also observed in a study that directly compared structural brain alterations in adult participants with schizophrenia and bipolar disorder^6^. Similarly, adults with schizophrenia exhibited widespread CT reductions^4^, while cortical thinning was restricted to frontal and temporal regions in bipolar disorder^5^. Both bipolar disorder and schizophrenia are associated with reduced volumes in the hippocampus and thalamus, but effects are of greater magnitude in schizophrenia^2,3^. Collectively, these findings suggest that widespread cortical SA and CT deficits are unique to schizophrenia, while subcortical structural abnormalities are present in both schizophrenia and bipolar disorder, albeit to a greater extent in schizophrenia.

Additionally, there are commonalities between the two disorders in terms of clinical characteristics, such as psychotic symptoms, and genetic risk variants^11,12^. For both disorders, medication effects may contribute to the observed brain alterations^4,5^. As such, it will be informative to investigate individuals at earlier stages of symptom emergence, prior to effects of chronic illness, as well as those who experience subclinical levels of symptomatology and thus are not medicated. Furthermore, many patients display mixed mood and psychotic symptoms^13^, yet it is not clear to what extent similar neuroanatomical alterations are observed in those who have both psychotic and bipolar spectrum symptoms compared to those with only psychosis or bipolar spectrum symptoms.

An initial investigation into volumetric abnormalities in youth with psychosis spectrum symptoms drawing from the same population sample utilized here, the Philadelphia Neurodevelopmental Cohort (PNC^14,15^), found that youth with psychosis spectrum symptoms exhibited reduced whole-brain gray matter, particularly in the medial temporal lobe and in frontal, temporal and parietal cortices^16^. Smaller scale studies of youth experiencing subclinical psychotic symptoms have found similar pattern of findings: reduced gray matter in frontal and temporal regions^17^. Furthermore, progressive medial orbitofrontal cortical thickness reductions have been observed in clinical high risk youth who convert to a psychotic disorder, indicating that these cortical thickness patterns may emerge prior to onset of psychosis^18^. However, the impact of mood symptoms, such as bipolar spectrum symptoms, on structural brain alterations has not yet been investigated. Furthermore, we do not know whether individuals who exhibit both psychosis and bipolar spectrum symptoms have distinct neuroanatomic alterations from these two groups.

To our knowledge, no studies have yet investigated the independent contributions of CT and SA to brain structure in psychosis and bipolar spectrum youth. Existing evidence shows that these indices are driven by different genetic and neurobiological mechanisms^8-10^. Furthermore, these subcomponents of brain volume offer a potentially meaningful window into the developmental course of brain structure. Early childhood brain development in healthy infants indicates cortical SA and CT develop separately, with cortical thickness achieving 97% of adult values on average by age two; in contrast, SA achieves only 69% of adult values by age two and continuing to further develop and expand^19^. As a result, these researchers suggested that SA explains most of cortical volume variation after two years, and proposed that early identification and prevention of neuropsychiatric illnesses focus on SA. However, this notion has not yet been fully explored, particularly with regards to youth who may be exhibiting early signs of serious mental illness.

As such, this study leveraged the PNC to examine cortical thickness, surface area, and subcortical volumes of psychosis spectrum (PS), bipolar spectrum (BP), both psychosis and bipolar spectrum (BP+PS), and typically developing (TD) youth, in order to determine: 1) common and distinct brain alterations in youth with psychosis and bipolar spectrum symptoms; 2) the stability of these deficits across development in comparison to TD youth; and 3) whether dimensional symptom severity is related to observed structural alterations. We hypothesized that: 1) PS youth would exhibit the most severe deficits, with the largest effect sizes occurring in prefrontal and temporal regions, and 2) within regions that exhibited group differences, PS youth would show the greatest age-associated alterations, relative to TD youth.

## Methods

### Sample

All data were obtained from the publicly available Philadelphia Neurodevelopmental Cohort (PNC, 1^st^ release, #7147) via the Database of Genotypes and Phenotypes (dbGap) platform. The data/analyses presented in the current publication are based on the use of study data downloaded from the dbGaP web site, under phs000607.v1.p1 (e.g., https://www.ncbi.nlm.nih.gov/projects/gap/cgi-bin/study.cgi?study_id=phs000607.v1.p1). The PNC is a population sample consisting of 9498 youth (ages 9-22 years) who participated in neurocognitive and genetic assessment after providing written informed consent or assent with parental consent (youth under 18 years old). A subset of these youth (N=997) also underwent neuroimaging. Study participants were assessed for psychiatric symptoms using the GOASSESS interview^20^, which incorporates questions from the Kiddie Schedule for Affective Disorders and Schizophrenia for School-Age Children (K-SADS^21^), the Structured Interview for Prodromal Syndromes (SIPS^22^), and the PRIME Screen Revised^23^. The TD group consisted of youth who denied clinically significant symptoms of psychopathology, based on responses to the GOASSESS interview. Similar to previous publications^16,20,24-26^, psychopathology was considered to be significant if symptoms endorsed were consisted with frequency and duration of a DSM-IV psychiatric disorder, while correspondingly accompanied by significant distress or impairment (a rating of ≥5 on a scale of 0-10). Individuals that endorsed symptoms meeting these criteria were excluded from the TD group, resulting in a final sample of N=376 youth.

The PS group was defined as in prior PNC publications^16,20,24-26^. Specifically, it included participants who: 1) had a score of 6 on any PRIME Screen Revised item; had a score of 5 or 6 on three or more items on the PRIME Screen Revised; or scored 2 standard deviations or more above the total score of age-cohort mean on the SIPS; or 2) answered yes to hallucination related questions on the KSADS and endorsed experiencing significant impairment or distress as a result, and not using drugs at the time of the experience of the symptom; or 3) scored 2 standard deviations or more above the age-cohort mean total score on 6 SOPS negative symptom items: attention and focus, disorganized speech, perception of self, experience of emotion, occupational function and avolition.

We defined a bipolar spectrum (BP) group that included participants who: 1) endorse at least two primary depressive symptoms on the KSADS, and 2) at least two primary manic or hypomanic symptoms, both lasting ≥1 day outside the context of substance use, illness, or medication use, with significant impairment or distress as a result of the symptoms.

### Clinical Symptoms & Functioning

Responses to the PS-R questionnaire were summed as a dimensional measure of positive symptoms. Responses were rated on a Likert scale (0=definitely disagree, 1=somewhat disagree, 2=slightly disagree, 3=not sure, 4=slightly agree, 5=somewhat agree, 6=definitely agree). Responses to mood-related symptoms on the KSADS (0=NO and 1=YES) were summed as a dimensional measure of mood. The exact items used for positive, negative, and mood symptoms are presented in Supplemental Table 1.

As part of the clinical testing, all participants were rated on the Global Assessment of Functioning (GAF), a widely used clinical scale to rate the social, occupational, and psychological functioning of an individual^27^. The scale ranges from 0-100, with higher scores indicating better overall functioning. We used current GAF scores as our measure of functioning.

### Neurocognitive Factor Scores

PNC participants underwent cognitive testing via the Penn Computerized Neurocognitive Battery (CNB). Descriptions of the metrics and calculation of efficiency scores for four domains were calculated according to a confirmatory factor analysis (N=9138^28^) for Complex Cognition (language reasoning, nonverbal reasoning and spatial ability), Executive Control (mental flexibility, attention, and working memory), Episodic Memory (verbal memory, face memory, and spatial memory) and Social Cognition (emotion identification, emotion differentiation, and age differentiation).

### Imaging processing and analysis

All neuroimaging data (N=997) were acquired on the same 3T Siemens TIM Trio MRI scanner at the Children’s Hospital at the University of Pennsylvania. The following imaging sequence was used for the T1-weighted images (MPRAGE): TR/TE/TI=1810/3.5/1000 ms, flip angle=9 degrees, slice thickness=1mm, FOV (RL/AP)=180/240 (as described in ^14^). The FreeSurfer image analysis suite (version 5.3.0) was used to derive bilateral measures of CT, cortical SA, and subcortical volume. FreeSurfer is a well-validated neuroimaging processing protocol that has previously been described in detail^29,30^. We extracted values based on the Desikan Freesurfer atlas^31^ and averaged the values from two hemispheres (34 regions for CT and SA; 6 subcortical volume regions). We implemented a quality assessment pipeline developed for the ENIGMA (Enhancing Neuroimaging Genetics through Meta-Analysis) consortium^32^ to assess individual scan quality. This pipeline has previously been implemented in multiple, large-scale studies of psychiatric disorders and of typical development ^3-5,33^.

### Statistical analyses

Data were analyzed using a mixed-effects regression with *lme4^34^* in R^35^. For all analyses, family membership (Supplemental Text) was included as a random effect and sex was included as a covariate. For regional SA, total SA of the estimated intracranial volume was included as a covariate. For subcortical measures, the total estimated intracranial volume was included as a covariate. For CT measures, overall mean CT was included as a covariate. We adjusted for these global measures because there was a significant main effect of group on all global sMRI measures (Supplementary Table 2). False discovery rate (FDR) correction was applied to p-values at a 0.05 level to control for multiple comparisons^36^. Equations for all analyses are presented in the Supplemental Text.

### Analysis 1: Group Differences in CT, SA, and subcortical volume

To determine an omnibus main effect across the four groups, we examined the overall main effects of group (typically developing youth (TD), PS, BP, and PS+BP) for each sMRI measure, while including age and the aforementioned variables as covariates. We followed up any significant main effects using the R package *lsmeans^37^* in order to investigate pairwise comparisons between groups.

### Analysis 2: Age-Associated Alterations in PS, BP, and/or PS+BP

We first examined linear, inverse, and quadratic forms of age within the typically developing group only. We used Aikaike’s information criterion (AIC), a commonly used measure for model selection^38^, to determine the model with the best fit (i.e., the lowest AIC). For each sMRI measure (i.e., each region of interest [ROI] examined), the inverse form of age (1/age) was considered to be the best fit. Then, to determine age-associated alterations that potentially differed between groups (PS, BP, PS+BP, TD), we examined an omnibus age by group interaction between the four groups for each sMRI measure. As in Analysis 1, we followed up any significant main effects by conducting pairwise comparisons of the least-squares means in each group (lsmeans, ^37^).

For any significant structural alterations identified in Analysis 1, we conducted an exploratory analysis by binning ages groups, using methods similar to what has previously used to characterize changes in late childhood (10_ 12 year-olds), early adolescence (13_15 year-olds), late adolescence (16-18 year-olds), and adulthood (19_22 year-olds). This parsing of groups is consistent with previous developmental publications (e.g., ^39,40^). After conducting a linear mixed model at each developmental stage for each region that significantly differed from controls, we estimated the R^2^ contribution of diagnosis using the R package *r2glmm^41^*.

### Analysis 3: Effects of Dimensional Measures

For any sMRI measure that exhibited a significant overall group difference in Analysis 1 or 2, we examined relationships with dimensional measures of symptoms and functioning, specifically: positive symptoms, negative symptoms, neurocognitive factor scores^28^ and current global functioning (GAF score). For any statistically significant relationships observed (*q*=0.05), we confirmed that this relationship remained present when group status was included as a factor in the model, in order to confirm that the relationship was still present once diagnosis status was taken into account. We also confirmed that there were no significant age by group interaction with any of the dimensional factors.

## Results

Participant information, including mean symptom measures and overall functioning scores, is presented in Table 1. Overall group differences in neurocognitive factor scores and corresponding pairwise contrasts are presented in Supplementary Table 2.

**Table 1.** Demographic and clinical information for all cohorts.

|  | TD<br>(N=376) | BP<br>(N=117) | BP+PS<br>(N=109) | PS<br>(N=113) |  |
| --- | --- | --- | --- | --- | --- |
| Age (mean years +/-SD) [range] | 15.1 +/-3.7 [9-22] | 16.8 +/- 3.5 [9-22]* | 15.9 +/- 3.0 [10-22] | 15.0 +/- 2.8 [9-22] | $p=1.5e-05$<br>BP>TD<br>BP>PS |
| Sex (M/F, % F) | 194/182, 48% | 56/61, 52% | 52/57, 52% | 57/56, 50% | $p=0.84$ |
| Mean Positive Symptoms (SD) | 2.5 (4.9) | 6.5 (7.2) | 24.5 (14.8) | 18.5 (12.1) | $p=2.2e-16$<br>BP+PS>BP<br>BP+PS>TD<br>BP+PS>PS<br>BP>TD<br>PS>BP<br>PS>TD |
| Mean Negative Symptoms (SD) | 1.0 (1.4) | 2.1 (2.0) | 4.3 (3.8) | 3.9 (3.7) | $p=2.2e-16$<br>BP+PS>BP<br>BP+PS>TD<br>BP>TD<br>PS>BP<br>PS>TD |
| Mean Depression Symptoms (SD) | 0.4 (1.1) | 4.1 (3.0) | 4.3 (2.9) | 1.2 (2.2) | $p=2.2e-16$<br>BP+PS>TD<br>BP+PS>PS<br>BP>TD<br>BP>PS<br>PS>TD |
| Mean Mania Symptoms (SD) | 0.7 (1.8) | 4.4 (2.8) | 6.0 (2.8) | 2.0 (2.9) | $p=2.2e-16$<br>BP+PS>BP<br>BP+PS>TD<br>BP+PS>PS<br>BP>PS<br>BP>PS<br>PS>TD |
| Current GAF (SD) | 85.0 (8.4) | 76.6 (11.8) | 72.3 (14.0) | 73.8 (11.9) | $p=2.2e-16$<br>TD>BP, BP+PS,<br>PS<br>BP>BP+PS |

Abbreviations are as follows: SD=standard deviation, M=Male, F=Female

### Global neuroanatomical measures

Typically developing youth had significantly larger estimated intracranial brain volumes (ICV) and larger total SA in comparison to all clinical groups (BP, PS, BP+PS). In comparison to the other three groups, BP+PS group exhibited significantly reduced overall mean CT (Supplementary Table 3, Supplementary Figure 1).

### Analysis 1: Reduced SA in Multiple Regions and Reduced Thalamic Volume are Specific to Psychosis Spectrum Youth

In comparison to all other groups (TD, BP, BP+PS), individuals in the PS group exhibited significantly reduced SA in lateral orbitofrontal, medial orbitofrontal, poster cingulate, precentral, and postcentral regions (Figure 1A-E, Table 2) as well as significantly reduced thalamic volume (Figure 1F, Table 3). BP+PS youth had greater medial orbitofrontal SA in comparison to TD and BP as well (Figure 1B). For all other sMRI measures, there were no significant pairwise differences between TD, BP, and BP+PS groups (Table 4).

**Table 2.** Overall statistics examining main effect of group (psychosis spectrum (PS), bipolar spectrum (BP), bipolar + psychosis spectrum (BP+PS), typically developing (TD)) on cortical thickness and surface area. For both measures, 1/Age, sex, and family membership were included in the model as covariates. For surface area measures, total surface area was included as an additional covariate. For cortical thickness measures, total mean cortical thickness was included as an additional covariate. Measures that passed statistical significance after correction for multiple comparisons (q<.05) are in bold.

| Brain Region | Cortical thickness |  |  | Surface Area |  |  |
| --- | --- | --- | --- | --- | --- | --- |
| | $\chi^2$ | $p$ | $q$ | $\chi^2$ | $p$ | $q$ |
| Banks of Superior Temporal Sulcus | 2.31 | 0.51 | 0.69 | 9.80 | 0.02 | 0.19 |
| Caudal Anterior Cingulate | 5.34 | 0.15 | 0.38 | 4.77 | 0.19 | 0.42 |
| Caudal Middle Frontal | 6.53 | 0.09 | 0.27 | 7.35 | 0.06 | 0.26 |
| Cuneus | 4.88 | 0.18 | 0.42 | 1.44 | 0.70 | 0.83 |
| Entorhinal | 0.59 | 0.90 | 0.92 | 2.14 | 0.54 | 0.72 |
| Fusiform | 1.65 | 0.65 | 0.81 | 6.80 | 0.08 | 0.26 |
| Inferior Parietal | 4.01 | 0.26 | 0.53 | 6.81 | 0.08 | 0.26 |
| Inferior Temporal | 0.28 | 0.96 | 0.96 | 4.18 | 0.24 | 0.52 |
| Isthmus Cingulate | 0.55 | 0.91 | 0.92 | 3.27 | 0.35 | 0.53 |
| Lateral Occipital | 7.75 | 0.05 | 0.26 | 3.42 | 0.33 | 0.53 |
| Lateral Orbitofrontal | 3.85 | 0.28 | 0.53 | **16.67** | **0.0008** | **0.02** |
| Lingual | 3.73 | 0.29 | 0.53 | 3.27 | 0.35 | 0.53 |
| Medial Orbitofrontal | 6.57 | 0.09 | 0.27 | **17.48** | **0.0006** | **0.02** |
| Middle Temporal | 0.86 | 0.83 | 0.89 | 8.47 | 0.04 | 0.25 |
| Parahippocampal | 5.02 | 0.17 | 0.41 | 3.32 | 0.35 | 0.53 |
| Paracentral | 0.60 | 0.90 | 0.92 | 3.25 | 0.35 | 0.53 |
| Pars Opercularis | 0.88 | 0.83 | 0.89 | 6.73 | 0.08 | 0.26 |
| Pars Orbitalis | 1.95 | 0.58 | 0.74 | 7.35 | 0.06 | 0.26 |
| Pars Triangularis | 1.24 | 0.74 | 0.85 | 7.26 | 0.06 | 0.26 |
| Pericalcarine | 2.30 | 0.51 | 0.69 | 2.29 | 0.52 | 0.69 |
| Postcentral | 9.20 | 0.03 | 0.20 | **13.49** | **0.004** | **0.05** |
| Posterior Cingulate | 2.05 | 0.56 | 0.73 | **13.18** | **0.004** | **0.05** |
| Precentral | 3.40 | 0.33 | 0.53 | **20.53** | **0.0001** | **0.01** |
| Precuneus | 3.38 | 0.34 | 0.53 | 4.73 | 0.19 | 0.42 |
| Rostral Anterior Cingulate | 2.84 | 0.42 | 0.60 | 6.05 | 0.11 | 0.32 |
| Rostral Middle Frontal | 5.50 | 0.14 | 0.37 | 5.86 | 0.12 | 0.33 |
| Superior Frontal | 3.68 | 0.30 | 0.53 | 7.54 | 0.06 | 0.26 |
| Superior Parietal | 1.34 | 0.72 | 0.84 | 9.51 | 0.02 | 0.19 |
| Superior Temporal | 3.92 | 0.27 | 0.53 | 3.44 | 0.33 | 0.53 |
| Supramarginal | 4.03 | 0.26 | 0.53 | 7.06 | 0.07 | 0.26 |
| Frontal Pole | 2.95 | 0.40 | 0.59 | 0.87 | 0.83 | 0.89 |
| Temporal Pole | 1.15 | 0.76 | 0.86 | 1.61 | 0.66 | 0.81 |
| Transverse Temporal | 3.61 | 0.31 | 0.53 | 10.08 | 0.02 | 0.19 |
| Insula | 0.61 | 0.89 | 0.92 | 7.10 | 0.07 | 0.26 |

**Table 3.** Overall statistics examining main effect of group (psychosis spectrum, bipolar spectrum, psychosis + bipolar spectrum, typically developing) on subcortical volume measures. 1/Age, sex, family membership, and total intracranial volume were all included in the model as covariates.

| Subcortical Structure | $\chi^2$ | *p* | *q* |
| --- | --- | --- | --- |
| Thalamus | **13.55** | **0.004** | **0.05** |
| Caudate | 1.44 | 0.70 | 0.75 |
| Putamen | 2.64 | 0.45 | 0.53 |
| Pallidum | 1.28 | 0.73 | 0.77 |
| Hippocampus | 5.24 | 0.16 | 0.27 |
| Amygdala | 7.75 | 0.05 | 0.17 |
| Nucleus Accumbens | 8.13 | 0.04 | 0.15 |

**Table 4.** Pairwise contrasts between groups for regions with a significant main effect of group (Tables 2 and 3).

| *Cortical Surface Area* | Contrast | Z-ratio | *p* |
| --- | --- | --- | --- |
| Lateral Orbitofrontal | TD > PS | 3.6 | 0.0003 |
|  | BP > PS | 2.3 | 0.02 |
|  | BP+PS>PS | 3.1 | 0.002 |
| Medial Orbitofrontal | TD > PS | 2.5 | 0.01 |
|  | BP+PS > PS | 3.9 | 0.0001 |
|  | BP+PS > BP | 2.6 | 0.009 |
|  | BP+PS > TD | 2.3 | 0.02 |
| Posterior Cingulate | TD > PS | 2.9 | 0.004 |
|  | BP > PS | 2 | 0.04 |
|  | BP+PS > PS | 3.4 | 0.0007 |
| Precentral Gyrus | TD > PS | 4.3 | 0.0001 |
|  | BP > PS | 2.6 | 0.009 |
|  | BP+PS > PS | 3.6 | 0.0003 |
| Postcentral Gyrus | TD > PS | 3.2 | 0.002 |
|  | BP > PS | 2.4 | 0.02 |
|  | BP+PS>PS | 3.2 | 0.001 |
| *Subcortical Volume* | Contrast | Z-ratio | *p* |
| Thalamus | TD > PS | 3.5 | 0.0005 |
|  | BP > PS | 2.1 | 0.04 |
|  | BP+PS > PS | 3 | 0.002 |

Overall group differences for all measures without global covariates (i.e., overall mean cortical thickness, total surface area, or estimated ICV) are reported in Supplementary Tables 4 and 5.

**Figure 1.**
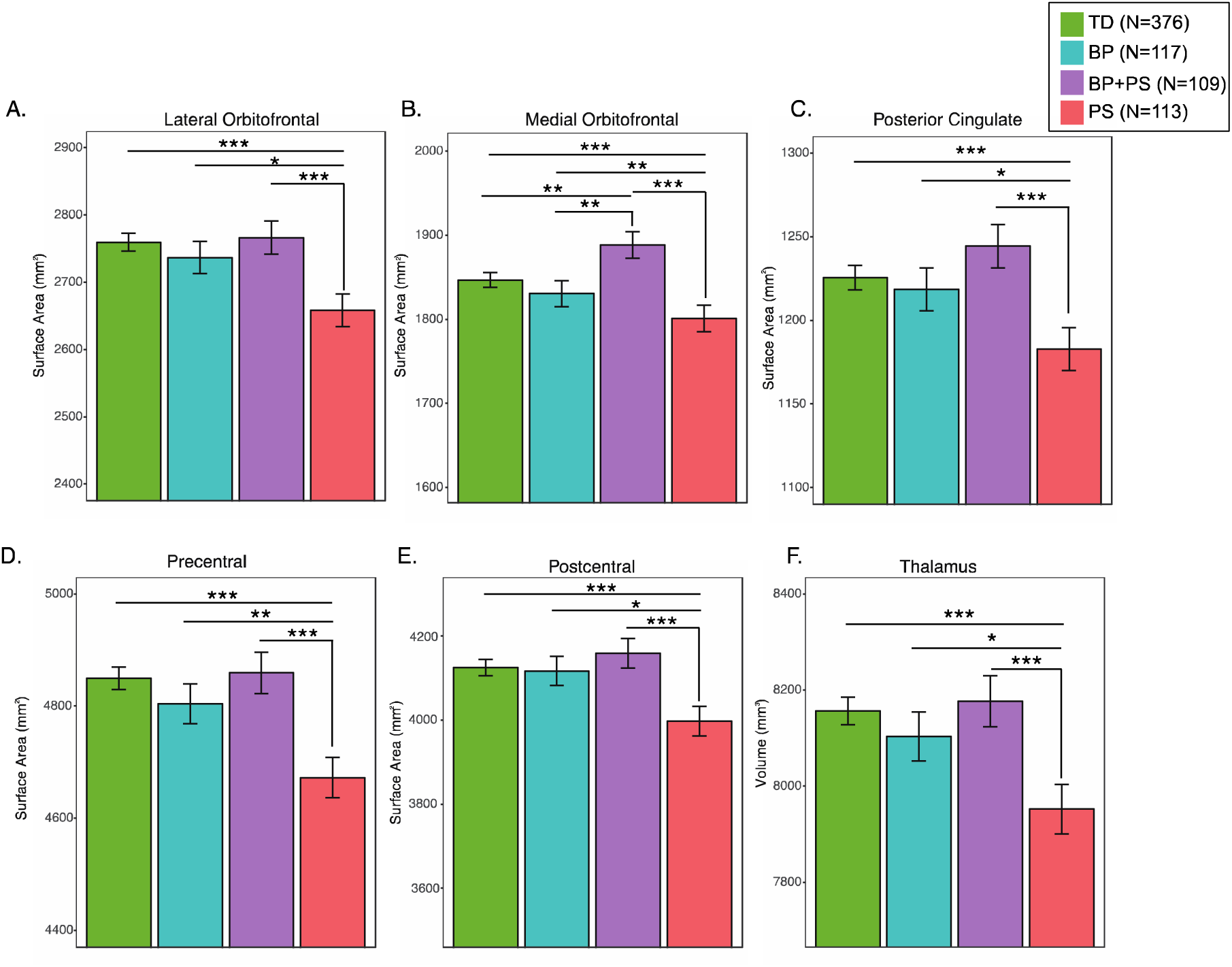
Psychosis spectrum (PS) youth exhibited reduced: A) lateral orbitofrontal, B) medial orbitofrontal, C) posterior cingulate, D) precentral, and E) postcentral surface area, in comparison to typically developing youth (TD), bipolar spectrum youth (BP), and youth with both BP+PS. \**p*<.05, \*\**p*<.01, \*\*\**p*<.005.

### Analysis 2: Structural alterations are greatest in early and late adolescence in psychosis spectrum youth

For all models, inverse age was the best fit (Supplementary Tables 6 and 7). Consistent with previous publications ^42-44^, 22 of the 34 regions exhibited cortical thinning with increasing age in TD youth (*q*<0.05). In this sample, largest effect sizes were observed for cingulate (isthmus and posterior) and temporal (middle and superior) regions. Eighteen out of 34 of the regions exhibited SA changes with age (q<.05). In the majority of these regions, SA decreased with increasing age. However, SA of the rostral anterior cingulate cortex increased with increasing age in TD (*q*=.01). In subcortical regions, TD exhibited increasing thalamic volume with increasing age(*q*=2e-6). The caudate, putamen, and palladium, and nucleus accumbens decreased in volume with increasing age (q<.05). However, across all groups, no significant age by group interaction remained statistically significant for any ROI measure (CT, SA, or subcortical volume) after FDR correction (Supplementary Tables 8 and 9).

Given that only the PS group exhibited neuroanatomical differences relative to TD, we focused our exploratory analyses on these two groups. Within separate developmental stages, we found that largest estimated effect size for group differences was during early adolescence for precentral and posterior cingulate regions, while largest estimated effect sizes for group differences in the thalamus, postcentral, and orbitofrontal regions were during late adolescence (Supplementary Table 10).

### Analysis 3: Surface Area is related to Cognition, Functioning, and Clinical Symptoms across Diagnostic Groups

Better functioning was associated with increased lateral orbitofrontal SA (*χ*^2^ =7.1, *p*=0.008, *q*=.02, Figure 2A). Higher complex cognition scores were associated with larger precentral SA (*χ*^2^ =19.0, *p*= 1.0e-5, *q*=.0001, Figure 2B). Higher executive functioning scale scores were associated with larger postcentral SA (*χ*^2^ =7.1, *p*= 0.008, *q*=.02, Figure 2C), while increased negative symptom severity was associated with reduced postcentral SA (%^2^=8.0, *p*=0.004, *q*=.02, Figure 2D). All associations remained statistically significant (*q*<0.05) when group status was included as factor in the model. There were no statistically significant relationships between neuroanatomic measures and positive symptoms of psychosis or mood symptoms (all *p*’s> 0.2).

Finally, post-hoc analyses showed that all results remained statistically significant when only one unique family member was included in the model.

**Figure 2.**
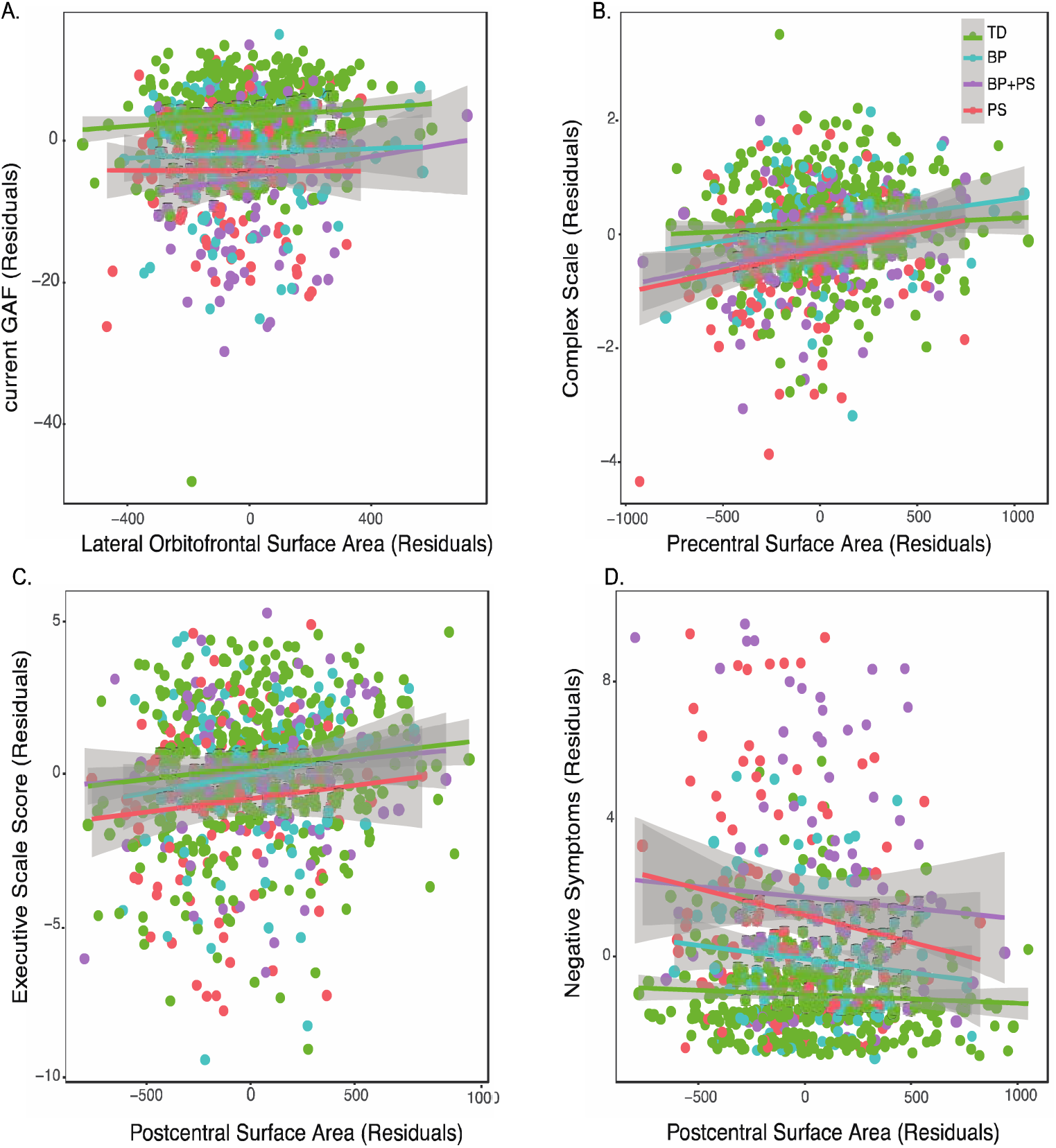
Across the entire sample: A) greater lateral orbitofrontal surface area was associated with higher global functioning scores, B) greater precentral surface was associated with higher Complex Cognition scores, C) greater postcentral surface area was associated with higher executive function scores, and D) reduced negative symptoms.

## Discussion

The goal of our study was to examine patterns of structural brain aberrations in youth experiencing subclinical symptoms of severe mental illness, to help us better understand abnormal developmental processes in distinct brain regions and determine more informative early biomarkers of illness. This analysis revealed several novel findings; specifically, we found that reduced surface area in multiple cortical regions (orbitofrontal, posterior cingulate, pre- and postcentral) was specific to psychosis spectrum (PS) youth in comparison to typically developing youth (TD), bipolar spectrum (BP) youth, and youth with both psychotic and bipolar symptoms (BP+PS). Among subcortical structure, we found reduced volume specifically in the thalamus in the PS group relative to these three groups. These findings provide support for the growing literature that neuroanatomic alterations are observable across the psychosis spectrum, and potentially prior to the onset of full-blown illness ^16,25^. Finally, we found that — across groups — higher overall functioning, less negative symptoms, and better higher-order cognition were associated with increased SA in some of these regions (specifically, the lateral orbitofrontal and postcentral regions), providing support for the notion that structural brain alterations reflect real-world behavior in youth. These dimensionally measured brain-behavior relationships highlight the importance of examining neural changes in a non-help seeking cohort, as our findings show that these neural alterations still have functional consequence, regardless of whether an individual develops a severe mental illness.

### Structural brain alterations are present in psychosis spectrum youth prior to adulthood

Consistent with multiple studies of youth at clinical high risk for developing psychosis ^45-47^, we found that youth with PS symptoms (***without*** co-occurring BP spectrum symptoms) exhibited reduced thalamic volume. Together, these findings suggest that reduced thalamic volumes are present in both help-seeking and non-help seeking youth with psychosis spectrum symptoms. Furthermore, these findings suggest that thalamic reductions, while prominent in schizophrenia ^48-50^, are not specific to the full-blown disorder and perhaps indicative of the extended psychosis phenotype.

In a prior study examining a larger cohort of PS youth (N=391), Satterthwaite et al ^16^ found that PS youth exhibited volumetric reductions in the posterior cingulate, orbitofrontal, and parietal cortices. We extend upon these findings, showing that the previously observed volumetric reductions in these regions are: 1) driven by SA reductions, and 2) that they are specific to PS youth who **do not** have concurrent bipolar spectrum symptoms. By examining separate components of volume (i.e., SA and CT) we can better understand the underlying neural dysfunction of PS symptoms. Specifically, converging evidence suggests that cortical SA is determined by proliferation of radial unit progenitors, which consist of neuroepithelial cells and radial glial cells^51^. Thus, the decreased SA observed in the orbitofrontal, precentral, postcentral, and posterior cingulate regions in PS youth may reflect reduced production of radial unit progenitors in these areas of the cortex. Given that, in this sample of TD youth, orbitofrontal SA was not significantly associated with age, this may be indicative of an early insult to brain development. Intriguingly, in youth with PS, SA differences are more circumscribed, and not as widespread as in adults with schizophrenia (e.g., ^4,6^). Furthermore, the effect sizes for SA when comparing TD to PS are stronger (Cohen’s d range 0.2-0.4) than the effect sizes identified in a recent multi-site consortium-wide meta-analysis comparing schizophrenia to healthy controls (Cohen’s d range: 0.05-0.1;^4^). These results suggest that SA reductions in orbitofrontal, pre- and postcentral, and posterior cingulate regions may reflect earliest risk markers of severe psychopathology and/or extended psychosis phenotype.

### Lack of CT Differences

Inconsistent with studies of adults with established schizophrenia and bipolar disorder diagnoses^4-6,52^, we did not identify statistically significant alterations in cortical thickness in youth with PS, BP, or BP+PS symptoms. The lack of findings could mean that cortical thinning observed in adults with schizophrenia and bipolar disorder reflects neuroanatomic changes that occur after the onset of overt psychiatric illness, due to effects of medications, and/or neurotoxic effects of chronic psychiatric illness. Alternatively, the absence of CT reductions in youth with subclinical PS and BP symptomatology could represent a marker of resilience.

### Unexpected patterns observed in psychosis + bipolar spectrum youth

Unexpectedly, we found structural alterations in youth with PS symptoms, but not in those with the combination of BP + PS symptoms. One proposed hypothesis is that the combination of mood and psychotic symptoms is associated with different underlying neurobiological mechanisms^53^. Is the experience of psychotic symptoms within the context of bipolar spectrum symptoms different than psychotic symptoms on their own, without prominent mood abnormalities? Intriguingly, individuals with both mood and psychotic symptoms have higher levels of functioning and better long-term outcome in comparison to those with schizophrenia, although these individuals are still impaired in comparison to healthy controls ^54,55^. In this sample, we found that BP + PS youth and PS youth had similar levels of global functioning. Thus, both groups are not functioning well, but show different biological patterns, suggesting that there may be different mechanisms driving this impairment.

### Developmental Implications

The age-associated cortical thinning observed in the TD in this sample is consistent with previous, longitudinal and cross-sectional studies of structural brain development during adolescence^42-44,56^. However, it is important to note that other longitudinal studies have identified different developmental trajectories of CT, including, non-linear CT changes from late childhood adolescence into adulthood (e.g.,^56-58^). Given that we studied a cross-sectional sample of TD, ability to detect non-linear changes in structural brain development are limited. Importantly, there is work showing that in a broader sample of PNC youth, implementation of general additive models identified non-linear age-associated changes in a few discrete brain regions^56^. However, this publication included youth that would have fallen into our PS, BP, or BP+PS groups. Given the importance of non-linear developmental changes in structural brain development, it will be important for future investigations to bring together multiple samples of typical development and utilize MRI harmonization methods to account for site effects ^59,60^ to allow for a more fine-grained analysis of age-associated disruption in youth at risk for and with psychiatric disorders.

In addition to identifying specific structural alterations in PS youth, we also observed preliminary evidence that the strongest effect sizes for group differences between typically developing youth and PS youth occurred during early and late adolescence. In regions that exhibited significant age associated changes in typical development (thalamic volume, posterior cingulate and postcentral SA), the period of early and late adolescence may be a particularly plastic or vulnerable stage in which an at-risk youth could “fall off’ of the normative developmental trajectory and be at greater risk for developing a psychiatric illness. Alternatively, these findings may be attributable to distinct developmental brain alterations, depending on the age at which one first begins to experience psychotic symptoms. It is important to note that this is an initial, descriptive analysis. Longitudinal studies of psychosis spectrum youth are necessary to probe these intriguing hypotheses. Growth charts, typically used as references for early identification of atypical development for metrics such a weight and head circumference^61^ have recently been extended to assess how psychiatric disorders are related to deviations from normative development^26,62^. In the future, multisite sample characterization of typical structural neurodevelopment will provide a template to further assess abnormal development of brain function in those with psychosis spectrum.

### Brain-Behavior Relationships

We found that greater lateral orbitofrontal surface area is related to global functioning, across groups. Functions associated with the lateral orbitofrontal cortex include evaluating possible outcomes based on contingencies and suppressing goal-irrelevant information to enable decision making or action^63-66^. It is plausible that lateral orbitofrontal structural aberrations contribute to impaired decision making, affecting one’s overall functioning. There is a wealth of evidence examining decision making in adults with an established diagnosis of schizophrenia and bipolar disorder^67-72^; however, the nature and extent of the relationship between brain maturation and the development of decision-making in youth, and across the broader psychosis spectrum, has yet to be explored. We also found that increased SA in the postcentral gyrus was associated with less severe negative symptoms and better executive cognition. The presence of this relationship suggests that reduced postcentral SA is a common underlying neural mechanism that contributes to both cognitive functioning and negative symptoms. Furthermore, given that these relationships were identified in the entire sample, these findings suggest that structural alterations in the postcentral region may be an important area for further study of transdiagnostic brain-behavior relationships.

### Limitations

This study was not without limitations, Importantly, we were not able to examine within-subject change over time in this cross-sectional sample. Furthermore, despite the large overall sample size, the number of individuals within each discrete developmental stage was more limited, particularly in the youngest and oldest groups. Given that age-associated changes can be small albeit meaningful, continued development of methods for combining data from different sites and scanners is warranted in order to fully map developmental changes in brain structure and function and their relevance to emerging psychiatric disorders. By examining a population-based, non-help-seeking sample, we were able to rule out possible confounds, such as illness chronicity and medication use; however, at the same time we do not know whether youth exhibiting PS and/or BP symptoms are destined to develop a full-blown major mental illness. However, the presence of subthreshold psychotic experiences (i.e., hallucinatory and delusional experiences) increases the likelihood that one will develop subsequent psychopathology^73-75^. Furthermore, though the majority of help-seeking youth deemed at risk for psychosis do not go on to develop overt psychosis, many continue to experience significant occupational and social impairment^76,77^. Thus, we may be tapping into identification of a cohort at risk for general psychopathology. These sorts of questions can only be answered with long term follow-up of participants as they mature.

### Conclusions & Future Directions

Taken together, our results provide compelling novel evidence for structural brain alterations specific to psychosis spectrum youth in adolescent neurodevelopment. These findings suggest potential biological distinctions between psychotic and bipolar spectrum conditions, which may suggest additional biomarkers relevant to early identification. Future, longitudinal studies focusing on typical ‘neurodevelopmental growth charts,’ and the extent and specific developmental periods during which youth at risk for serious mental illnesses deviate from these trajectories, are necessary to address many remaining questions regarding brain alterations relevant to emergent psychopathology in vulnerable youth.

## Acknowledgements

We would like to acknowledge the following funding sources: R01 MH107250 Genetic Risk for Developmental Expression of Neuropsychiatric Intermediate Traits (Carrie E. Bearden & Roel A. Ophoff), K01 MH112774 Neurodevelopmental Variation of Intrinsic Functional Connectivity and Its Relationship to Psychosis Risk and Gene Expression (Maria Jalbrzikowski), RC2 MH089983 Neurodevelopmental Genomics: Trajectories of Complex Phenotypes (Raquel E. Gur).

## Presentation Information

This study was presented as an abstract at the International Congress on Schizophrenia Research Meeting, March 24-28 2017, San Diego, CA.

